# Varied influence of curli on Shiga toxin-producing *Escherichia coli* O157, O26, O111 biofilm formation and animal cell adherence

**DOI:** 10.64898/2026.09.15.751941

**Authors:** Indira T. Kudva, Erika N. Biernbaum, Hannah Mazon, Koy Drabek

## Abstract

Shiga toxin-producing *Escherichia coli* (STEC) persist in cattle and can colonize the bovine recto-anal junction, making it an important reservoir for transmission. Curli are extracellular amyloid fibers associated with biofilm formation and environmental persistence, but their contribution to STEC adherence to bovine epithelial cells is unclear. This study examined the relationship between curli production, biofilm formation, and adherence of O157, O26, and O111 STEC isolates to bovine recto-anal junction (RAJ) squamous epithelial (RSE) cells. Thirty isolates were evaluated for RSE adherence, curli-associated phenotypes, and biofilm formation under environmental and host-associated growth conditions. O157 and O111 isolates predominantly exhibited strong aggregative adherence to RSE cells, whereas O26 isolates showed more heterogeneous, primarily diffuse adherence. O111 isolates produced the strongest biofilms under environmental conditions and displayed relatively stable curli phenotypes, while O157 and O26 isolates showed greater variation. However, environmental biofilm formation did not consistently correlate with RSE adherence. Biofilm formation was also markedly reduced under host-simulated conditions in DMEM-LG at 26, 37, and 39°C, including among isolates that produced strong environmental biofilms. Differences in Shiga toxin genotype and expression likewise did not account for the major adherence patterns, as O111 isolates lacking *stx_2_* retained strong aggregative adherence. These findings indicate that curli-associated biofilm formation is primarily influenced by environmental conditions and does not directly predict STEC adherence to bovine RSE cells. The results suggest that environmental persistence and host-cell attachment represent distinct phenotypes and that STEC adherence to bovine epithelial cells is likely mediated by additional strain- and serotype-specific factors.

**IMPORTANCE:** Our results highlight a paradoxical role for curli in STEC biofilm formation versus host cell adherence that has implications in both diagnostics and intervention strategies. Standard laboratory biofilm assays may not accurately reflect true STEC-host cell adherence phenotypes, and curli targeted for environmental sanitation may not reduce persistence at the bovine RAJ suggesting a need for multi-target pre-harvest interventions. A complex interplay of curli and non-curli factors, besides growth conditions, appears to influence environmental versus bovine RSE cell attachment that do not always correlate with genotypes.

## INTRODUCTION

Shiga toxin-producing *Escherichia coli* (STEC) are important foodborne pathogens that colonize the gastrointestinal tract of healthy cattle, the principal reservoir for human infection (1, 2). While *E. coli* O157:H7 has historically been the predominant focus of STEC research, non-O157 serogroups are increasingly recognized as significant causes of human disease and are frequently isolated from cattle and beef products worldwide (3–5). Successful colonization of cattle depends on the ability of STEC to adhere to epithelial surfaces within the recto-anal junction (RAJ), a preferred site of bacterial persistence and shedding (6). Adherence to recto-anal squamous epithelial (RSE) cells is considered a critical early event in colonization, facilitating long-term carriage and increasing the likelihood of environmental dissemination and contamination of the food chain (7). Consequently, identifying bacterial factors that promote adherence to the bovine RAJ is essential for developing intervention strategies aimed at reducing STEC carriage in cattle.

Among the surface structures implicated in STEC persistence, curli fimbriae have emerged as multifunctional amyloid fibers that contribute to adhesion, biofilm formation, environmental survival, and interactions with host tissues. Curli are major structural components of the extracellular matrix in *E. coli* biofilms and promote attachment to both biotic and abiotic surfaces, enhancing bacterial resistance to environmental stresses encountered in agricultural and food-processing settings (8, 9). Numerous studies have demonstrated that both O157 and non-O157 STEC vary considerably in their ability to produce biofilms, with curli expression representing one of several factors associated with robust biofilm development and persistence (9–12). Curli-producing STEC have also been associated with enhanced attachment to food-contact surfaces and increased tolerance to sanitizers, suggesting that these fibers facilitate persistence throughout the farm-to-food continuum (10, 12).

In addition to their role in environmental persistence, accumulating evidence indicates that curli contribute directly to host colonization. Curli have been shown to promote MAC-T epithelial cell invasion, biofilm formation, and persistence of *E. coli* O157:H7 in cattle (13). We previously demonstrated that curli tempers adherence of O157:H7 to bovine RSE cells in a strain-dependent manner, indicating that the contribution of curli to host colonization is more complex than a simple adhesin-mediated interaction (14). However, whether curli play a comparable role in adherence by the diverse non-O157 STEC serotypes remains unknown. Given the substantial genetic and phenotypic diversity among STEC and the reported variability in curli production and biofilm-forming capacity, it cannot be assumed that observations made with O157:H7 apply broadly across non-O157 strains (9, 11, 15).

The present study extends our previous investigation of O157:H7 by evaluating the role of curli in adherence to bovine RSE cells across a broader collection of O157 and non-O157 STEC isolates. Because curli are well established as determinants of biofilm formation and environmental persistence, determining whether they similarly influence colonization of the bovine RAJ may reveal common mechanisms underlying both environmental survival and host persistence. Such knowledge will improve our understanding of STEC biology in cattle and may identify conserved targets for interventions designed to reduce colonization, fecal shedding, and subsequent transmission through the food production chain.

## RESULTS

### STEC isolate diversity and serotype-specific clustering was observed by DNA fingerprinting

DNA fingerprinting *via* PATS PCR separated the strains by serotype and all of the isolates were diverse in terms of their individual profiles (Tables 1 and S1). Notably, several of the polymorphic *Xba*I sites (IK8, IK114, IK118, IK123, and IK127) were absent in all STEC O26 and O111 isolates (apart from IK123 in O111-10) but mostly present in the STEC O157 isolates, and IK25 was absent in all 30 isolates (Table S1). The only *Xba*I sites present in the STEC O26 isolates were IKB3 and IKB5, with IKB5 present in 8/10 STEC O111 isolates. There were intact IKNR7, IKNR10, and IKNR12 *Avr*II restriction sites in 28/30 of the isolates, while IKNR3 and IKNR33 were either present with a SNP or intact in most STEC O157 but absent in all STEC O26 and O111 isolates. Additionally, *Avr*II sites IKNR16 and IKNR27 were intact in STEC O157 but contained a SNP in STEC O26 and O111 isolates. Shiga toxin gene distribution varied among isolates, as all STEC O157 had *stx2* but was absent in most STEC O26, while all STEC O111 had *stx1*. All 30 STEC O157, O26, and O111 isolates possessed *eaeA* and *hlyA*.

Additionally, the thirty STEC isolates could be grouped based on their PATS patterns, producing 17 unique profiles: 8 profiles for the O157 isolates, 4 for the O26 isolates, and 5 for the O111 isolates (Table 1). Distribution of isolate origin was also somewhat represented by profile grouping: type III were of either ground beef or bovine origin; types IV, V, XIII, XVI, and XIX were of human origin; types VII, VIII, IX, and X were of ground beef origin; types VI and XI were of bovine origin; human and bovine origins in types XII, XV, and XVII; and fly and human origins for types XIV and XVIII (Tables 1, 3 and S1).

**Table 1.** Summary of PATS pattern types.

| PAT<br>S<br>Type | Polymorphic<sup>1</sup> *Xba*I sites |  |  |  |  |  |  |  | Polymorphic<sup>2</sup> *Avr*II sites |  |  |  |  |  |  | Virulence<br>Genes |  |  |  | Strain |
| --- | --- | --- | --- | --- | --- | --- | --- | --- | --- | --- | --- | --- | --- | --- | --- | --- | --- | --- | --- | --- |
|  | IK8 | IK25 | IK114 | IK118 | IK123 | IK127 | IKB3 | IKB5 | IKNR3 | IKNR7 | IKNR10 | IKNR12 | IKNR16 | IKNR27 | IKNR33 | *stx*<sub>1</sub> | *stx*<sub>2</sub> | *eaeA* | *hlyA* |  |
| C<sup>3</sup> | 1 | 0 | 1 | 1 | 1 | 1 | 1 | 0 | 2 | 2 | 2 | 2 | 1 | 2 | 2 | 0 | 1 | 1 | 1 | STEC O157 strain RM6067W |
| C<sup>3</sup> | 0 | 0 | 0 | 0 | 0 | 1 | 0 | 0 | 0 | 0 | 0 | 0 | 1 | 1 | 0 | 0 | 0 | 0 | 0 | *E. coli* strain K-12 |
| C<sup>3</sup> &<br>I | 0 | 0 | 1 | 1 | 1 | 1 | 1 | 1 | 2 | 2 | 2 | 2 | 2 | 2 | 2 | 1 | 1 | 1 | 1 | STEC O157 strain EDL933(ATCC 43895) and<br>O157-9 |
| II | 1 | 0 | 1 | 1 | 1 | 1 | 0 | 0 | 2 | 1 | 1 | 2 | 1 | 2 | 2 | 0 | 1 | 1 | 1 | O157-1 |
| III | 1 | 0 | 1 | 1 | 1 | 1 | 0 | 1 | 2 | 2 | 2 | 2 | 2 | 2 | 2 | 1 | 1 | 1 | 1 | O157-2 and O157-3 |
| IV | 0 | 0 | 1 | 1 | 1 | 0 | 0 | 0 | 0 | 2 | 2 | 2 | 1 | 2 | 2 | 0 | 1 | 1 | 1 | O157-4 |
| V | 1 | 0 | 0 | 1 | 1 | 1 | 0 | 0 | 1 | 2 | 2 | 2 | 1 | 2 | 0 | 0 | 1 | 1 | 1 | O157-5 and O157-6 |
| VI | 1 | 0 | 1 | 1 | 1 | 1 | 1 | 1 | 2 | 2 | 2 | 2 | 2 | 2 | 2 | 1 | 1 | 1 | 1 | O157-7 |
| VII | 1 | 0 | 0 | 1 | 1 | 1 | 1 | 0 | 1 | 2 | 2 | 2 | 1 | 2 | 2 | 1 | 1 | 1 | 1 | O157-8 |
| VIII | 1 | 0 | 0 | 1 | 1 | 1 | 1 | 0 | 1 | 2 | 2 | 2 | 1 | 2 | 0 | 1 | 1 | 1 | 1 | O157-10 |
| IX | 0 | 0 | 0 | 0 | 0 | 0 | 1 | 1 | 0 | 0 | 0 | 0 | 1 | 1 | 0 | 0 | 1 | 1 | 1 | O26-1 |
| X | 0 | 0 | 0 | 0 | 0 | 0 | 0 | 0 | 0 | 2 | 2 | 2 | 1 | 1 | 0 | 1 | 0 | 1 | 1 | O26-2, O26-4, O26-6, O26-8, O26-9, and O26-10 |
| XI | 0 | 0 | 0 | 0 | 0 | 0 | 1 | 0 | 0 | 2 | 2 | 2 | 1 | 1 | 0 | 1 | 1 | 1 | 1 | O26-3 |
| XII | 0 | 0 | 0 | 0 | 0 | 0 | 1 | 0 | 0 | 2 | 2 | 2 | 1 | 1 | 0 | 1 | 0 | 1 | 1 | O26-5 and O26-7 |
| XIII | 0 | 0 | 0 | 0 | 0 | 0 | 0 | 1 | 0 | 2 | 2 | 2 | 1 | 1 | 0 | 1 | 0 | 1 | 1 | O111-1 and O111-6 |
| XIV | 0 | 0 | 0 | 0 | 0 | 0 | 0 | 0 | 0 | 2 | 2 | 2 | 1 | 1 | 0 | 1 | 1 | 1 | 1 | O111-3 |
| XV | 0 | 0 | 0 | 0 | 0 | 0 | 0 | 1 | 0 | 2 | 2 | 2 | 1 | 1 | 0 | 1 | 1 | 1 | 1 | O111-4 and O111-5 |
| XVI | 0 | 0 | 0 | 0 | 0 | 0 | 1 | 1 | 0 | 2 | 2 | 2 | 1 | 1 | 0 | 1 | 1 | 1 | 1 | O111-2, O111-7, O111-8, and O111-9 |
| XVII | 0 | 0 | 0 | 0 | 1 | 0 | 0 | 0 | 0 | 2 | 2 | 2 | 1 | 1 | 0 | 1 | 0 | 1 | 1 | O111-10 |

<sup>1</sup>0, absence of amplicon; 1, presence of amplicon.
<sup>2</sup>1, the *Avr*II restriction site has a small nucleotide polymorphism (SNP); 2, the *Avr*II restriction site is intact
<sup>3</sup>C, control

**Table 2.** EHEC-ELISA absorbance values used to score Shiga toxin expression by STEC isolates.

| Strain | Average Post-ELISA Absorbance (450nm) | Standard Deviation | Toxin Gene<sup>1</sup> / Expression Score<sup>2</sup> | *stx*<sub>1</sub> | *stx*<sub>2</sub> |
| --- | --- | --- | --- | --- | --- |
| Kit + control | 1.571 | ± 0.02416 | n/a<sup>3</sup> | n/a | n/a |
| Kit - control | 0.078 | ± 0.001886 | n/a | n/a | n/a |
| C<sup>4</sup>: STEC O157 strain RM6067W | 3.145 | ± 0.1442 | 2 / +++ | - | + |
| C<sup>4</sup>: STEC O157 strain EDL933 (ATCC 43895) | 2.668 | ± 0.002239 | 3 / +++ | + | + |
| C<sup>4</sup>: *E. coli* strain K-12 | 0.09067 | ± 0.01155 | - | - | - |
| O157-1 | 2.079 | ± 0.01202 | 2 / ++ | - | + |
| O157-2 | 2.737 | ± 0.338 | 3 / +++ | + | + |
| O157-3 | 3.377 | ± 0.3613 | 3 / +++ | + | + |
| O157-4 | 1.181 | ± 0.08415 | 2 / + | - | + |
| O157-5 | 0.584 | ± 0.03606 | 2 / + | - | + |
| O157-6 | 0.655 | ± 0.1838 | 2 / + | - | + |
| O157-7 | 2.152 | ± 0.6477 | 3 / ++ | + | + |
| O157-8 | 3.14 | ± 0.3755 | 3 / +++ | + | + |
| O157-9 | 2.372 | ± 0.2934 | 3 / ++ | + | + |
| O157-10 | 2.756 | ± 0.08768 | 3 / +++ | + | + |
| O26-1 | 0.644 | ± 0.009899 | 2 / + | - | + |
| O26-2 | 2.432 | ± 0.5459 | 1 / ++ | + | - |
| O26-3 | 3.015 | ± 0.01202 | 3 / +++ | + | + |
| O26-4 | 3.105 | ± 0.02263 | 1 / +++ | + | - |
| O26-5 | 3.061 | ± 0.1075 | 1 / +++ | + | - |
| O26-6 | 2.486 | ± 0.09405 | 1 / ++ | + | - |
| O26-7 | 3.127 | ± 0.4165 | 1 / +++ | + | - |
| O26-8 | 3.013 | $\pm 0.8648$ | 1 / +++ | + | - |
| O26-9 | 3.034 | $\pm 0.2616$ | 1 / +++ | + | - |
| O26-10 | 3.045 | $\pm 0.384$ | 1 / +++ | + | - |
| O111-1 | 3.189 | $\pm 0.08273$ | 1 / +++ | + | - |
| O111-2 | 1.83 | $\pm 0.00495$ | 3 / ++ | + | + |
| O111-3 | 3.041 | $\pm 0.1138$ | 3 / +++ | + | + |
| O111-4 | 2.924 | $\pm 0.6746$ | 3 / +++ | + | + |
| O111-5 | 3.223 | $\pm 0.06293$ | 3 / +++ | + | + |
| O111-6 | 3.086 | $\pm 0.09051$ | 1 / +++ | + | - |
| O111-7 | 2.389 | $\pm 0.08273$ | 3 / ++ | + | + |
| O111-8 | 2.36 | $\pm 0.0601$ | 3 / ++ | + | + |
| O111-9 | 3.37 | $\pm 0.2369$ | 3 / +++ | + | + |
| O111-10 | 3.017 | $\pm 0.07$ | 1 / +++ | + | - |

<sup>1</sup>Toxin gene score: 1, *stx*<sub>1</sub> only; 2, *stx*<sub>2</sub> only; 3, *stx*<sub>1</sub>+ *stx*<sub>2</sub>
<sup>2</sup>Toxin expression score: +, post assay *A*<sub>450</sub> is 0.19 - 1.5; ++, 1.6 - 2.5; +++, 2.6 - 3.5
<sup>3</sup>n/a, not applicable
<sup>4</sup>C, control

**Table 3.** Strains used in this study.

| No. | Strain ID<sup>1</sup> | Origin | Virulence Genes |
| --- | --- | --- | --- |
| **O157:H7 strains** |  |  |  |
| 1 | USDA 2 | Human | *stx*<sub>1</sub><sup>-</sup>, *stx*<sub>2</sub><sup>+</sup>, *eae*<sup>+</sup>, *hlyA*<sup>+</sup> |
| 2 | USDA 39 | Human | *stx*<sub>1</sub><sup>+</sup>, *stx*<sub>2</sub><sup>+</sup>, *eae*<sup>+</sup>, *hlyA*<sup>+</sup> |
| 3 | USDA 49 (RM6049) | Human | *stx*<sub>1</sub><sup>+</sup>, *stx*<sub>2</sub><sup>+</sup>, *eae*<sup>+</sup>, *hlyA*<sup>+</sup> |
| 4 | TX 909-1 | Bovine | *stx*<sub>1</sub><sup>-</sup>, *stx*<sub>2</sub><sup>+</sup>, *eae*<sup>+</sup>, *hlyA*<sup>+</sup> |
| 5 | FSIS-10 | Ground beef | *stx*<sub>1</sub><sup>-</sup>, *stx*<sub>2</sub><sup>+</sup>, *eae*<sup>+</sup>, *hlyA*<sup>+</sup> |
| 6 | FSIS-11 | Ground beef | *stx*<sub>1</sub><sup>-</sup>, *stx*<sub>2</sub><sup>+</sup>, *eae*<sup>+</sup>, *hlyA*<sup>+</sup> |
| 7 | FSIS-53 | Ground beef | *stx*<sub>1</sub><sup>+</sup>, *stx*<sub>2</sub><sup>+</sup>, *eae*<sup>+</sup>, *hlyA*<sup>+</sup> |
| 8 | FSIS-62 | Ground beef | *stx*<sub>1</sub><sup>+</sup>, *stx*<sub>2</sub><sup>+</sup>, *eae*<sup>+</sup>, *hlyA*<sup>+</sup> |
| 9 | FSIS-71 | Ground beef | *stx*<sub>1</sub><sup>+</sup>, *stx*<sub>2</sub><sup>+</sup>, *eae*<sup>+</sup>, *hlyA*<sup>+</sup> |
| 10 | FSIS-78 | Ground beef | *stx*<sub>1</sub><sup>+</sup>, *stx*<sub>2</sub><sup>+</sup>, *eae*<sup>+</sup>, *hlyA*<sup>+</sup> |
| **O26:H11 strains** |  |  |  |
| 1 | 7-14 50A | Bovine | *stx*<sub>1</sub><sup>-</sup>, *stx*<sub>2</sub><sup>+</sup>, *eae*<sup>+</sup>, *hlyA*<sup>+</sup> |
| 2 | DECA-10A | Human | *stx*<sub>1</sub><sup>+</sup>, *stx*<sub>2</sub><sup>-</sup>, *eae*<sup>+</sup>, *hlyA*<sup>+</sup> |
| 3 | 10205 | Human | *stx*<sub>1</sub><sup>+</sup>, *stx*<sub>2</sub><sup>+</sup>, *eae*<sup>+</sup>, *hlyA*<sup>+</sup> |
| 4 | 99.0703 | Bovine | *stx*<sub>1</sub><sup>+</sup>, *stx*<sub>2</sub><sup>-</sup>, *eae*<sup>+</sup>, *hlyA*<sup>+</sup> |
| 5 | EHEC2-TW04272 | Human | *stx*<sub>1</sub><sup>+</sup>, *stx*<sub>2</sub><sup>-</sup>, *eae*<sup>+</sup>, *hlyA*<sup>+</sup> |
| 6 | P5-1 | Bovine | *stx*<sub>1</sub><sup>+</sup>, *stx*<sub>2</sub><sup>-</sup>, *eae*<sup>+</sup>, *hlyA*<sup>+</sup> |
| 7 | EHEC2-CL5 | Not confirmed | *stx*<sub>1</sub><sup>+</sup>, *stx*<sub>2</sub><sup>-</sup>, *eae*<sup>+</sup>, *hlyA*<sup>+</sup> |
| 8 | 18 B24-2 | Bovine | *stx*<sub>1</sub><sup>+</sup>, *stx*<sub>2</sub><sup>-</sup>, *eae*<sup>+</sup>, *hlyA*<sup>+</sup> |
| 9 | DEC-10B | Human | *stx*<sub>1</sub><sup>+</sup>, *stx*<sub>2</sub><sup>-</sup>, *eae*<sup>+</sup>, *hlyA*<sup>+</sup> |
| 10 | DEC-10E | Bovine | *stx*<sub>1</sub><sup>+</sup>, *stx*<sub>2</sub><sup>-</sup>, *eae*<sup>+</sup>, *hlyA*<sup>+</sup> |
| **O111:H8 strains** |  |  |  |
| 1 | 7-14 10A | Bovine | *stx*<sub>1</sub><sup>+</sup>, *stx*<sub>2</sub><sup>-</sup>, *eae*<sup>+</sup>, *hlyA*<sup>+</sup> |
| 2 | 1056-1 | Bovine | *stx*<sub>1</sub><sup>+</sup>, *stx*<sub>2</sub><sup>+</sup>, *eae*<sup>+</sup>, *hlyA*<sup>+</sup> |
| 3 | DECA-8B | Human | *stx*<sub>1</sub><sup>+</sup>, *stx*<sub>2</sub><sup>+</sup>, *eae*<sup>+</sup>, *hlyA*<sup>+</sup> |
| 4 | 90.0102 | Bovine | *stx*<sub>1</sub><sup>+</sup>, *stx*<sub>2</sub><sup>+</sup>, *eae*<sup>+</sup>, *hlyA*<sup>+</sup> |
| 5 | IDPH14614 | Human | *stx*<sub>1</sub><sup>+</sup>, *stx*<sub>2</sub><sup>+</sup>, *eae*<sup>+</sup>, *hlyA*<sup>+</sup> |
| 6 | DEC-8A | Human | *stx*<sub>1</sub><sup>+</sup>, *stx*<sub>2</sub><sup>-</sup>, *eae*<sup>+</sup>, *hlyA*<sup>+</sup> |
| 7 | 7-1664A | Fly | *stx*<sub>1</sub><sup>+</sup>, *stx*<sub>2</sub><sup>+</sup>, *eae*<sup>+</sup>, *hlyA*<sup>+</sup> |
| 8 | 7-4580A | Fly | *stx*<sub>1</sub><sup>+</sup>, *stx*<sub>2</sub><sup>+</sup>, *eae*<sup>+</sup>, *hlyA*<sup>+</sup> |
| 9 | EHEC2-TW05614 | Human | *stx*<sub>1</sub><sup>+</sup>, *stx*<sub>2</sub><sup>+</sup>, *eae*<sup>+</sup>, *hlyA*<sup>+</sup> |
| 10 | BEO0-749 | Human | *stx*<sub>1</sub><sup>+</sup>, *stx*<sub>2</sub><sup>-</sup>, *eae*<sup>+</sup>, *hlyA*<sup>+</sup> |

<sup>1</sup>(10)

### STEC O157 isolates were the most diverse in Shiga toxin expression

An EHEC-ELISA was conducted to confirm and quantitate Shiga toxin expression among the 30 isolates. As the Premier® EHEC kit does not differentiate between *stx_1_* and *stx_2_*, we coupled the ELISA with PCR to produce a toxin gene / expression score. Isolates without *stx_1_* had a lower A_450_ reading than those with *stx_1_* (O157-1, O157-4, O157-5, O157-6, and O26-1; Fig 1). The average post-ELISA absorbance among most STEC O26 and O111 isolates was very similar (Table 2). The majority of STEC O111 isolates (7/10) possessed both Shiga toxins. Most of the STEC O26 isolates (8/10) had only *stx_1_*, while no STEC O157 isolate was *stx_1_*-only. Interestingly, *stx_2_* was present in all ground beef isolates.

**Figure 1.**
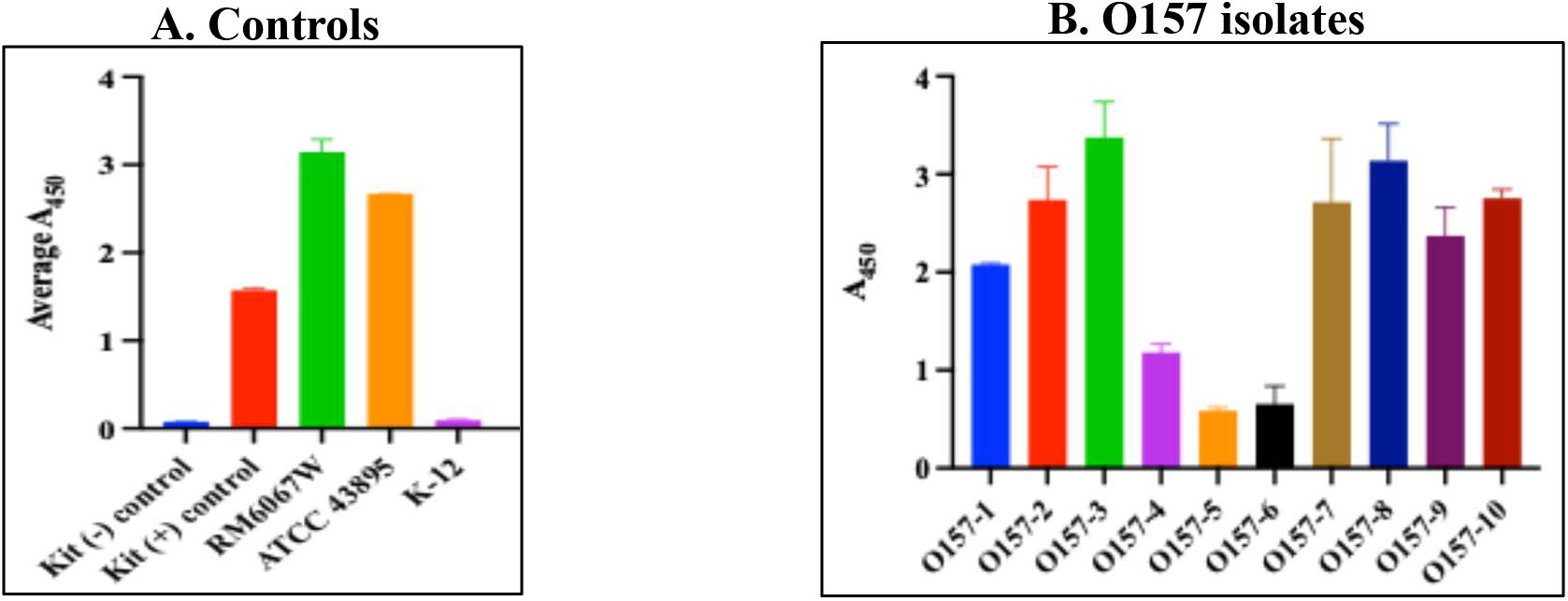
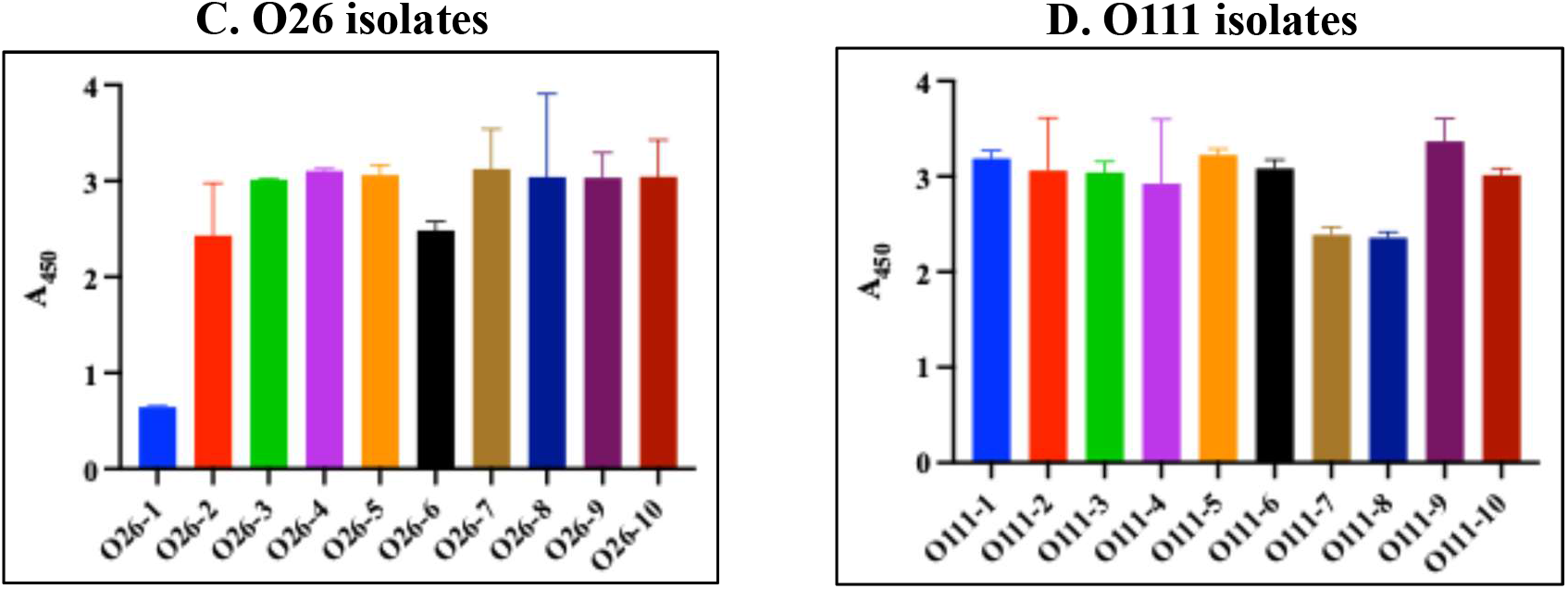
Shiga toxin expression among STEC O157, O26, and O111 isolates. Average post-ELISA A_450nm_ readings with standard deviation are displayed along the y-axis and isolates along the x-axis. Panel A: assay controls of Premier® kit positive- and negative-controls, STEC O157 strains RM6067W and EDL933/ATCC 43895, and *E. coli* strain K-12/K-12. Panels B-D: STEC O157 isolates (B), O26 isolates (C), or O111 isolates (D).

Toxin gene / expression scores were based upon both the distribution of Shiga toxin genes and the quantity of gene expression among isolates. As seen in Table 2, there was a variation of scores within serotypes. O157 isolates were diverse with scores fairly evenly divided between“3 / +++” and “2 / +” with a couple of isolates categorized as “3 / ++” and one as “2 / ++”. O26 isolates consisted primarily of “1 / +++”, with two “1 / ++”, and one each of “2 / +” and “3 / +++”. Lastly, O111 isolates were the least diverse of the serotypes, consisting only of “3 / +++”, “3 / ++”, and “1 / +++” toxin gene / expression scores. In all, the *stx* gene was functional in all 30 isolates.

### STEC O157 and O111 isolates demonstrated the strongest adherence to the bovine rectoanal junction squamous epithelial (RSE) cells

Adherence patterns of the thirty STEC isolates to RSE cells were evaluated in an *in vitro* assay, as previously reported (14, 16, 17). Despite being dis-similar serotypes with distinct PATS profiles and variations in Shiga toxin phenotypes, the STEC O157 and STEC O111 isolates shared the same aggregative RSE adherence pattern with most of the RSE cells having >10 bacteria attached (Figs. 2A, C and S1; Table S2). Analysis of the quantitative adherence data (Table S2) yielded no significant differences between the isolates within each of these two serotypes. A peripheral role in attachment for Stx2 (18), predominantly present in the STEC O157 and O111isolates, could not be supported by our observations as the STEC O111 isolates without this toxin gene also demonstrated aggregative adherence.

**Figure 2.**
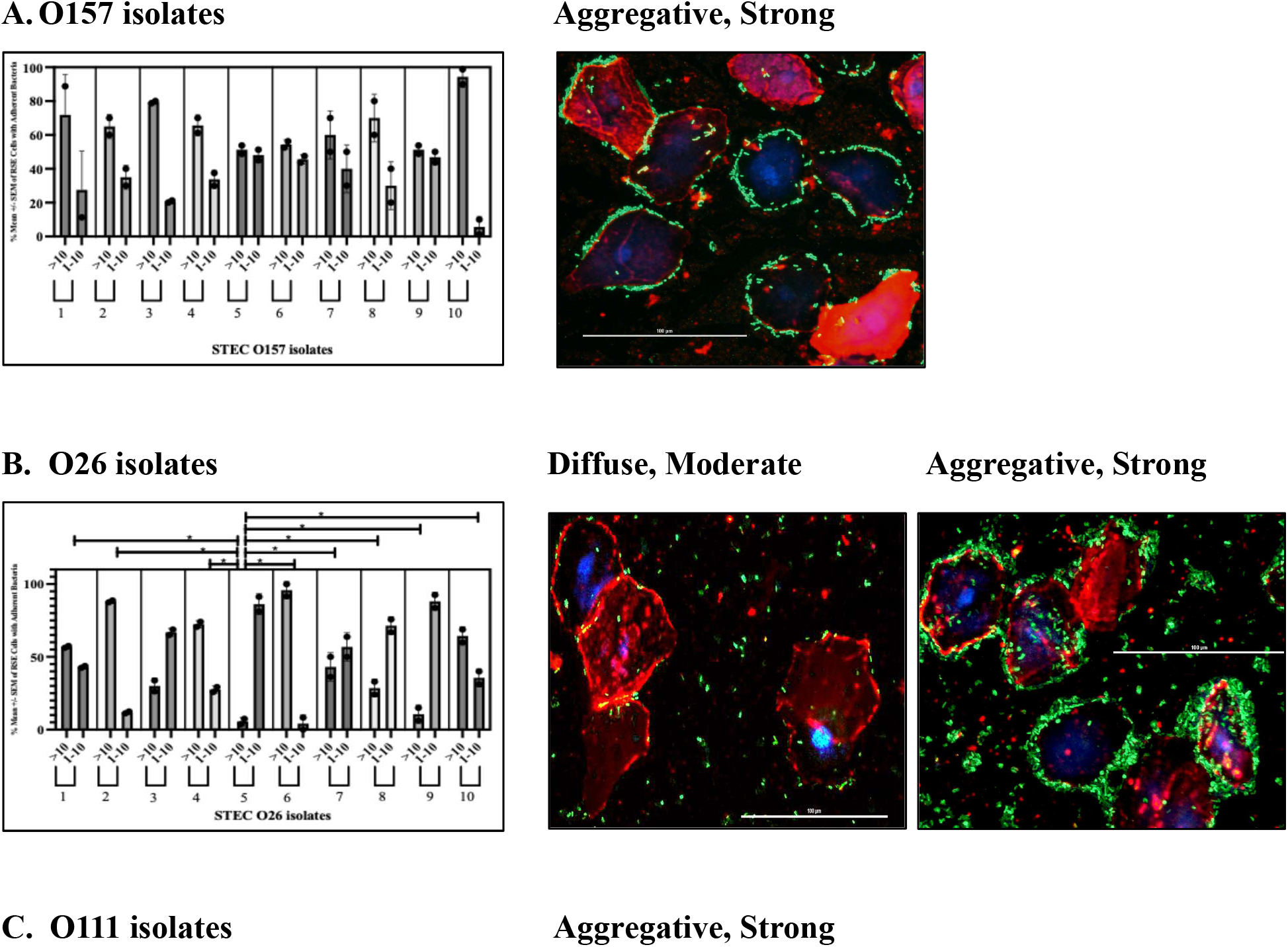
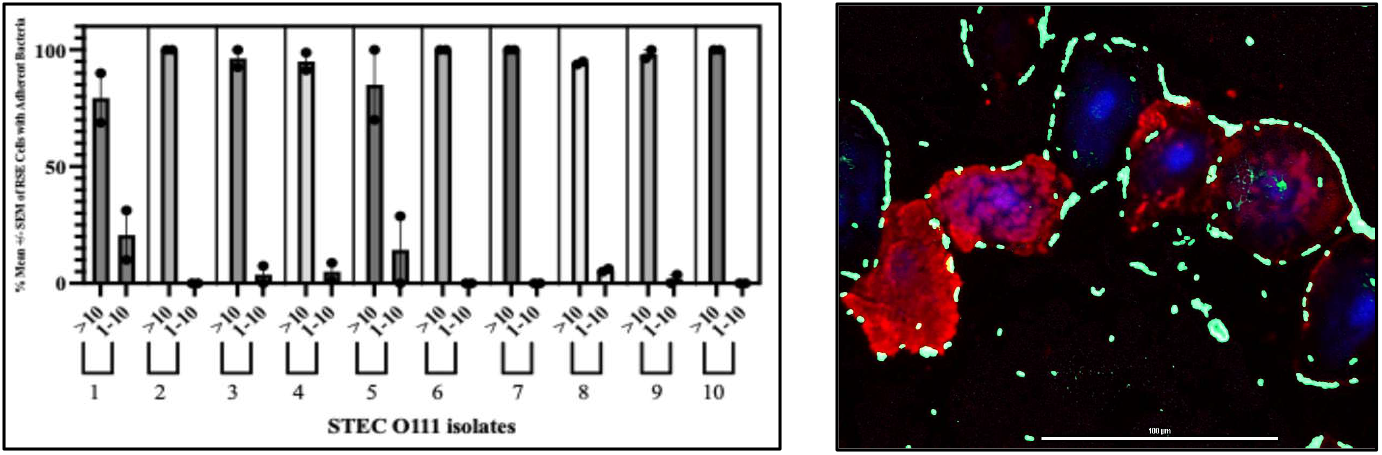
Quantitative and qualitative representation of STEC adherence on RSE cells. Graphs represent percent mean ± standard error of mean for number of RSE cells with >10 and 1-10 adherent bacteria per isolate within each serotype. Details of actual counts are in Table S2; each of two assays conducted had one slide per treatment group. Each slide in turn had two sets of 4 technical replicates spotted on it in two rows and 20 well-dispersed cells were evaluated for adherent bacteria per spot per row. The accompanying immunofluorescence-stained images are shown at 40x magnification and represent the common adherence phenotypes observed per serotype. Bacteria have green fluorescence, RSE cells’ cytokeratins have orange-red fluorescence, and the nuclei have blue fluorescence.

The STEC O26 isolates on the other hand, demonstrated contrasting diffuse adherence patterns except for one isolate #O26-6 that had an aggregative adherence on the RSE cells (Figs. 2B and S1; Table S2). The diverse nature of attachment resulted in statistically significant quantitative differences among the STEC O26 isolates as shown in the graph (Fig 2B); the *p-* values were, *p* = 0.0117 for each of the O26 isolates 1, 2, 4, 6, 7, 8, 10 versus O26-5, and *p* = 0.0345 between O26-5 and O26-9. Control strains produced expected phenotypes of aggregative adherence for STEC O157 strains RM6067W and EDL933, and diffuse adherence for *E. coli* K-12 (Fig S1).

### STEC O111 were strong biofilm producers under environmental growth conditions and demonstrated a steady, non-variant curli phenotype

All thirty isolates were initially evaluated for biofilm formation in 2-day plate assays in LB-NS at 26°C and sub-cultured on CRI agar plates. Curli-positive but biofilm-negative isolates were further evaluated in 5-day tube assays. If an isolate had a mixed CRI phenotype, the different colony types were isolated and tested separately under similar conditions as above. The phenotypes on CRI plates and biofilms were scored as shown in Table S3.

Five of the ten STEC O157 isolates were curli- and biofilm-negative (#1, 4-6, 10), and among the remaining isolates, #2, 3, 7, 8 had mixed curli phenotypes and biofilm formation while O157-9 produced pink colonies and delayed biofilm formation (Table S3; Fig 3). Of note, isolate O157-8 did not form a smooth suspension when grown in LB-NS at both 26°C and 37°C but was smooth in both DMEM-LG and DMEM-NG. The red colonies from this isolate (O157-8R), along with those of O157-7 (O157-7R), also displayed a red-dry-rough (rdar) phenotype on CRI agar. These two isolates also produced the most biofilm of the ten O157 isolates and the respective variants that were evaluated (Fig 4). Six of the STEC O26 isolates were biofilm-negative (#2, 3, 5-7, 9), O26-8 was positive for biofilm formation with pink colonies on CRI plates, while O26-1, O26-4, and O26-10 displayed mixed curli and biofilm phenotypes (all of bovine origin).

**Figure 3.**
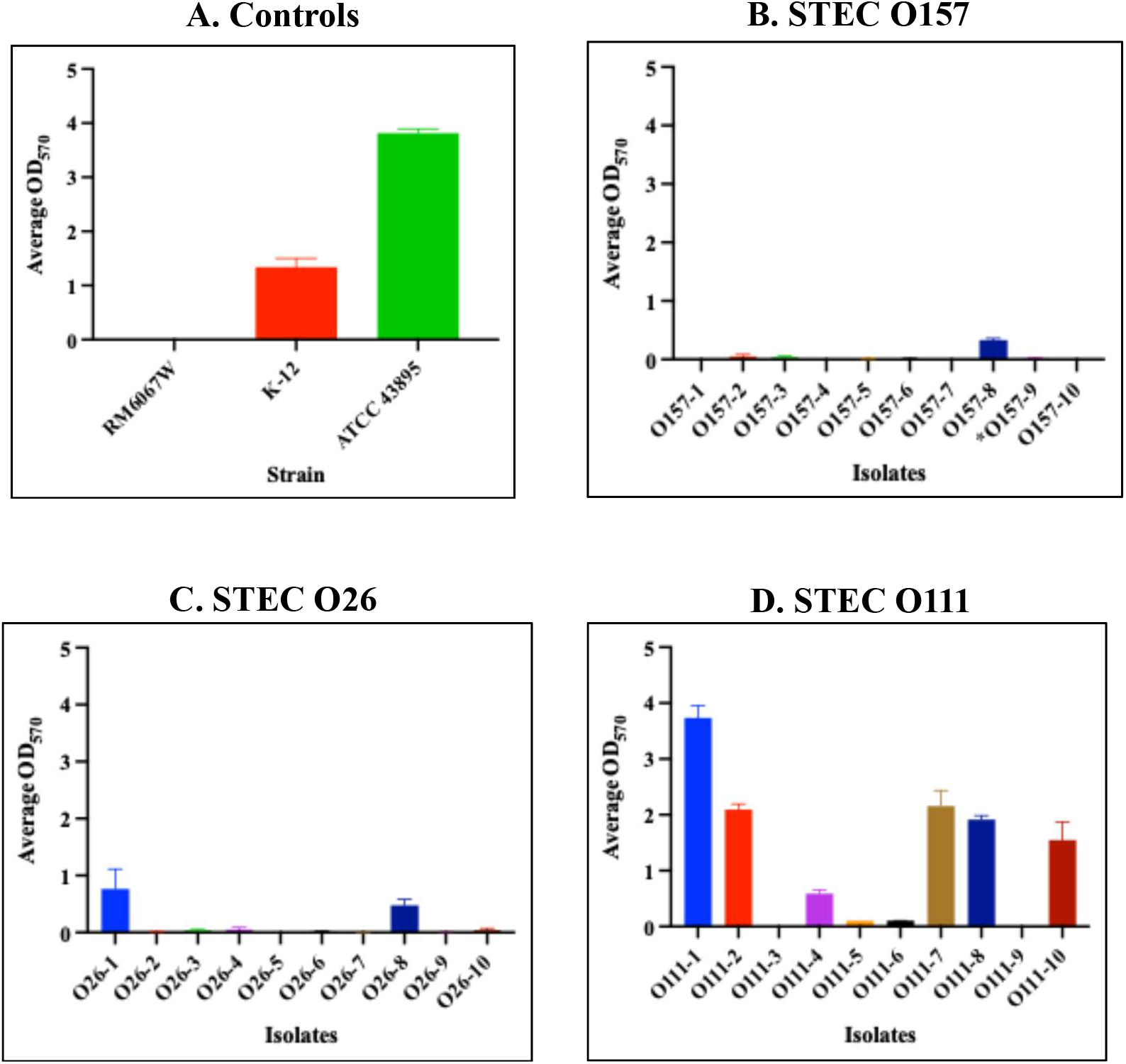
Biofilm graphs of the 30 STEC isolates. Assays were conducted in LB-NS and microtiter plates were incubated for 2 d at 26°C. Average post-solubilization OD_570nm_ readings with standard deviation are displayed along the y-axis and isolates along the x-axis. Panel A: assay control strains; panels B-D: STEC O157 isolates (B), STEC O26 isolates (C), or STEC O111 isolates (D).

**Figure 4.**
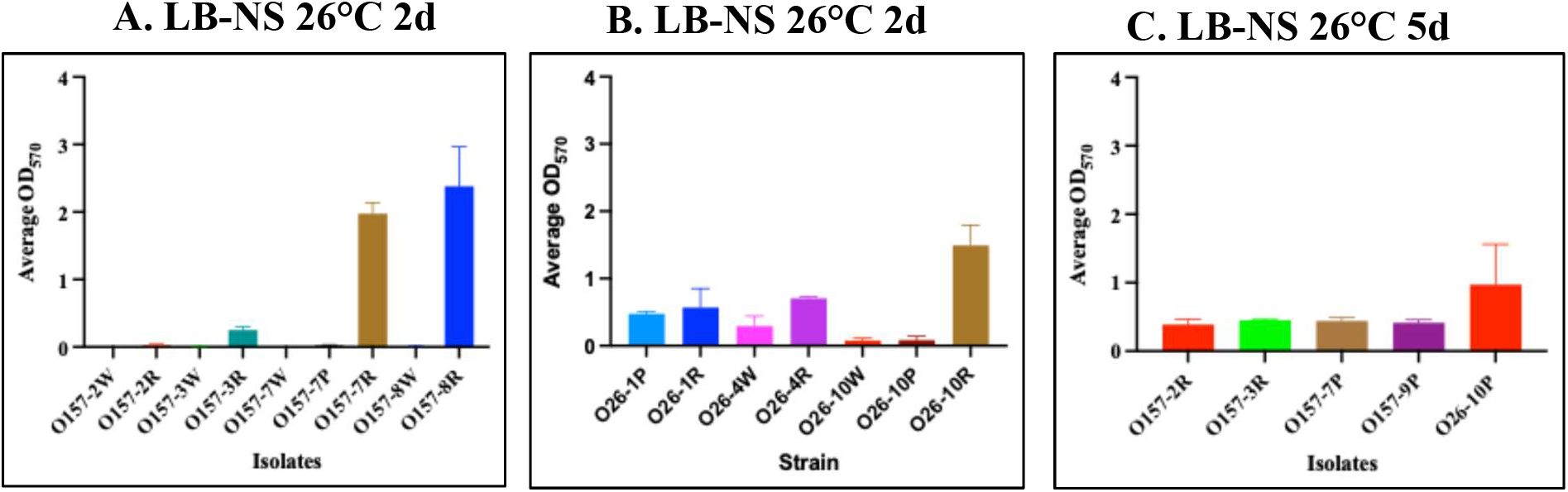
Biofilm graphs of isolated red-white curli variants. Mixed CRI phenotypes were first isolated and evaluated separately in LB-NS at 26°C. Assays included both 2 d microtiter plate and 5 d tube, of which the average post-solubilization OD_570nm_ readings with standard deviation are displayed along the y-axis and isolates along the x-axis. Panel A: O157 red-white isolates in a 2 d plate assay; Panel B: O26 red-white isolates in a 2 d plate assay; Panel C: O157 and O26 red-white isolates in a 5 d tube assay.

Interestingly, none of the STEC O111 isolates produced mixed curli phenotypes, as they were either curli positive (#1, 2, 4, 7, 8, 10) or curli negative (#3, 5, 6, 9). O111-1 produced a biofilm similar to that of O157 strain EDL933/ATCC 43895 and was the only isolate of the thirty to do so, with O111-7 producing slightly less. Biofilm production for O157-7R, O26-10R, O111-2, O111-8, and O111-10 was similar to *E.coli* strain K-12 (Figs 3 and Fig 4). Furthermore, O157-2R, O157-3R, O157-7P, O157-9P, and O26-10P appear to be slow biofilm formers as a weak biofilm was only observed for these isolates after 5 days of incubation at 26°C. The STEC O111 isolates had the highest biofilm production, with O157 producing the least. Additionally, the bovine isolates among all 30 strains produced the most biofilm, followed by the two fly isolates, while the majority of the human and ground beef isolates were non-biofilm producers.

### Immunofluorescent staining verified absence of curli expression in some STEC isolates with the white phenotype on CRI plates

Seventeen STEC isolates with a definite white phenotype on the CRI plates were evaluated for curli production *via* IF staining for curlin, along with the control strains (Fig S2). These included the STEC O157 isolates# 1W, 2W, 3W, 4W, 5W, 6W, 7W, 8W, 10W; O26 isolates# 2W, 5W, 9W, 10W; and O111 isolates# 3W, 5W, 6W, 9W. Following a 5 d tube assay for biofilms, planktonic and biofilm forming bacteria were separately stained for curlin. As expected, biofilms were negligible for most test isolates when compared to the controls O157 stain EDL933/ATCC43895 and *E. coli* K-12.

As shown in Fig S2, the majority of the isolates that produced white colonies on the CRI plates did not produce curli; in such instances, both planktonic and weak biofilm-associated bacteria were not stained by the antibodies targeting curlin. The eight exceptions were, O157-4W, 5W, 6W, 7W, 8W, 10W; O26-9W, and O111-6W that had few bacteria still expressing curli especially in the planktonic growth (Fig S2). Presence of even the few curli-expressing bacteria could be indicative that curli presence/absence may not be simply structural but more regulatory and is not uniform across the bacterial population.

### STEC produce weak or no biofilms under host-simulated growth conditions

The strong biofilms produced by STEC O111 under environmental conditions could be extrapolated to the aggregative RSE cell adherence demonstrated by isolates of this serotype (Fig 2C). However, a similar association could not be made with any of the isolates of STEC O157 or isolate O26-6 that also demonstrated aggregative RSE cell adherence but made weak or no biofilms in LB-NS at 26°C (Fig 2A-B).

Hence, the control strains and a few STEC isolates with varied curli phenotype on CRI plates at 26°C (Table S3; EDL933, O157-1W, O157-9P, O26-6P, O26-9W, O26-10R, O111-2R, O111-3W and O111-8R), derived from parent strains with different RSE cell adherence patterns (Table S2), were further tested in 2 d biofilm assays with DMEM-LG at 26°C, 37°C and 39°C while comparing against the 2 d assay in LB-NS at 26°C. This was done to ascertain any differences in biofilm formation under human/bovine host-simulated (DMEM-LG at 37°C or 39°C) versus environmental conditions and verify plausible role for curli in RSE cell adherence.

Negligible-to-no biofilms were produced by the control strains and test isolates in DMEM-LG at 26°C, 37°C and 39°C (Fig 5, Table S4). *E. coli* K-12, STEC O157 strain EDL933, O111-2R and O111-8R that made strong biofilms in LB-NS at 26°C made poor biofilms in DMEM-LG at all temperatures (Fig 5B-D); STEC O157 strain EDL933 created a slightly better biofilm compared to other isolates tested at 39°C and to the strain itself at 26°C or 37°C (Fig 5D, Table S4). These results suggest a minimal or peripheral role for curli in STEC attachment to host cells, especially bovine intestinal cells, unlike with biofilm formation in the environment.

**Figure 5.**
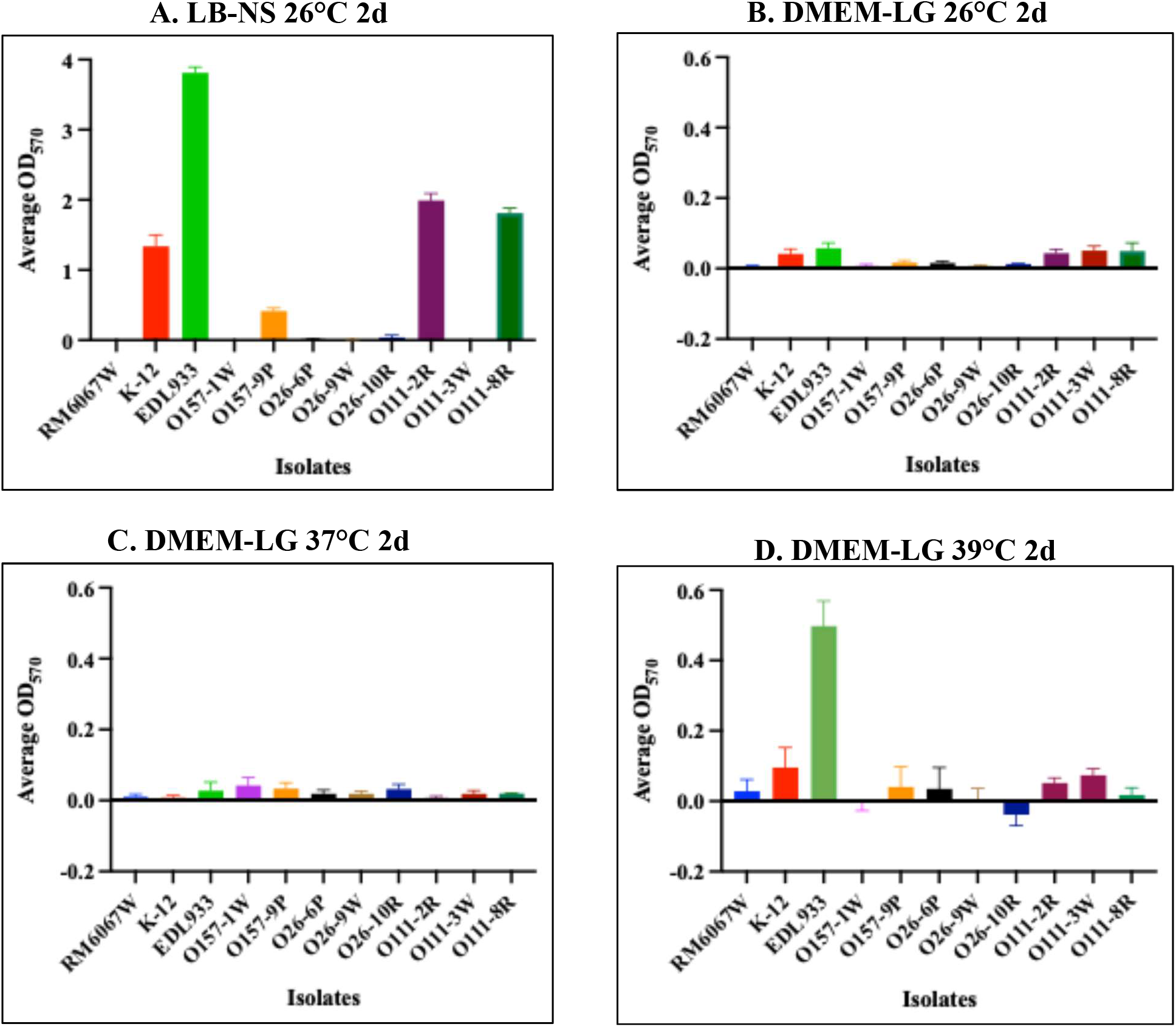
Biofilm graphs of select STEC isolates with varied curli and adherence phenotypes. Isolates were initially evaluated in LB-NS at 26°C for 2 d (panel A) and subsequently in DMEM-LG for 2 d at either 26°C (panel B), 37°C (panel C) or 39°C (panel D). The average post-solubilization OD_570nm_ readings with standard deviation are displayed along the y-axis and isolates along the x-axis. To accommodate the negligible levels of biofilm formed in DMEM-LG at all incubation temperatures, lower OD_570nm_ ranges are depicted on the y-axis for graphs in panels B-D.

### Host-simulated growth conditions appeared to impact curli expression in some of the STEC isolates

Since the tested STEC isolates and control strains produced weak-to-no biofilms under host-simulated growth conditions, these bacteria were cultured on CRI plates using the same growth conditions as for the biofilm assay. This was to verify if curli expression was negatively impacted by growth in LB-NS broth at 26°C versus DMEM-LG broth, at 37°C and 39°C.

Only three of the tested STEC isolates, O26-10R, O111-2R and O111-8R displayed different CRI phenotypes post-growth in DMEM-LG at 37°C and 39°C, than in LB-NS at 26°C (Fig S3). O26-10R produced red colonies on CRI plates following growth in LB-NS broth at 26°C but had a mixed red and white phenotype after growth in DMEM-LG at 37°C and 39°C. O111-2R was smooth, pink in DMEM-LG at 37°C and 39°C versus red after growth in LB-NS at 26°C (Fig S3). Similarly, O111-8R was white in DMEM-LG at 37°C and 39°C versus red after growth in LB-NS at 26°C (Fig S3). All control strains and other tested STEC isolates maintained their phenotypes on the CRI plates irrespective of the incubation temperatures or broth used to initiate cultures (Fig S3).

## DISCUSSION

The objective of this study was to investigate whether curli, a surface-associated structure strongly implicated in environmental persistence and biofilm formation by *Escherichia coli*, contributes to adherence of STEC to bovine recto-anal junction squamous epithelial (RSE) cells. The results demonstrate that the relationship between curli production, biofilm formation, and host-cell adherence is complex and appears to depend strongly on bacterial serotype and environmental conditions. Although substantial variation in curli-associated phenotypes was observed among the STEC isolates, the strongest RSE adherence was observed among O157 and O111 isolates irrespective of their curli phenotype. However, although the O111 isolates exhibited the strongest biofilm-forming capacity under environmental conditions, this phenotype was markedly reduced when bacteria were grown in host-associated medium. Collectively, these findings suggest that the mechanisms supporting environmental biofilm formation and adherence to bovine epithelial cells are not necessarily similar and that curli may have a more limited or indirect role in STEC colonization of the bovine recto-anal junction.

The RSE assay was originally developed specifically to examine bacterial factors involved in attachment to the bovine recto-anal junction, an important site of STEC persistence in cattle (19). The O157 and O111 isolates demonstrated a predominantly aggregative and strong adherence phenotype on bovine RSE cells, with most RSE cells having >10 adherent bacteria. In contrast, the O26 isolates were considerably more heterogeneous, with most displaying diffuse adherence and only O26-6 exhibiting a strong aggregative phenotype. These observations are consistent with previous studies demonstrating that STEC adherence to bovine RSE cells is strain- and serotype-dependent with different mechanisms, distinct from those used to attach to human epithelial cells, responsible for adherence of non-O157 versus O157 STEC (17, 20). The contrasting O26 phenotype further supports the possibility that multiple, serotype-or strain-specific adherence mechanisms function at the bovine epithelial cell surface. The observation that O26-6 exhibited aggregative adherence while the remaining O26 isolates were predominantly diffuse further indicates that serotype alone does not determine the adherence phenotype. Recent comparative genomic work examining *E. coli* adherence to bovine RSE cells similarly identified several accessory gene clusters associated with high adherence, although none of the associations reached statistical significance (21). These findings emphasize the likelihood that RSE adherence is multifactorial and may involve previously uncharacterized bacterial surface factors (21).

A clear relationship between curli expression and environmental biofilm formation was observed, particularly among the O111 isolates. Under LB-no-salt (LB-NS) conditions at 26°C, STEC O111 isolates were the strongest biofilm producers, whereas O157 isolates generally produced less biofilm and several O157 and O26 isolates were biofilm negative. The O111 population also differed from O157 and O26 in exhibiting relatively stable curli phenotypes rather than the mixed red/white phenotypes observed among several O157 and O26 isolates. These findings are consistent with the established role of curli as an important component of the extracellular matrix of *E. coli* biofilms that is under strong environmental regulation (8, 9). Leech et al. demonstrated that medium composition, temperature, osmolarity, and other environmental parameters substantially influence biofilm development and curli gene expression in *E. coli* (22). Thus, the strong biofilm phenotype observed in LB-no-salt at 26°C likely represents an environmental persistence phenotype rather than a direct measure for the ability to attach to bovine epithelial cells.

The heterogeneous curli phenotype observed among the O157 isolates is also consistent with previous work demonstrating considerable variation in curli expression among O157 strains. Uhlich et al. demonstrated substantial differences in curli expression, including stable red and white phenotypes in otherwise closely related O157 variants (23). Red variants exhibited approximately fourfold greater *csgD* promoter activity than white variants, illustrating how regulatory variation can produce pronounced phenotypic differences in curli production. Similarly, the occurrence of mixed curli phenotypes in the present set of isolates may reflect varied regulation rather than simply the absence or presence of the *csg* genes. This interpretation is supported by our immunofluorescence observations. Although many isolates with a white phenotype on Congo red indicator plates did not show detectable curli, a subset retained a small population of curli-expressing cells. Thus, the present results suggest that curli expression may be heterogeneous within individual STEC populations. Such phenotypic heterogeneity could be biologically advantageous because it allows bacteria to simultaneously maintain subpopulations adapted to environmental persistence and other subpopulations likely better suited to host-associated growth (24).

The most important finding of this study is that strong environmental biofilm formation did not consistently correspond to strong RSE-cell adherence. The O111 isolates provide the clearest example for this. These isolates were the strongest biofilm producers under environmental conditions and also exhibited strong aggregative RSE adherence. However, a direct causal relationship between these two phenotypes could not be inferred because strong RSE adherence was also observed among O157 isolates and O26-6 despite their weak or absent biofilm formation under the same environmental conditions. More importantly, when selected isolates were evaluated under host-associated conditions using DMEM-LG, biofilm formation was reduced to negligible or low levels at 26°C, 37°C, and 39°C. Strains such as *E. coli* K-12, O157 EDL933, O111-2R, and O111-8R that formed substantial biofilms in LB-no-salt at 26°C showed substantially weaker biofilm formation in DMEM-LG; only EDL933 showed a biofilm phenotype at 39°C. Thus, biofilm formation appeared to be highly dependent on the growth environment/conditions. Overall, these findings indicate that environmental curli-dependent biofilm formation is not predictive of STEC’s ability to adhere to bovine epithelial cells. The distinction is biologically plausible because the bacterium encounters very different physicochemical conditions in an environmental reservoir compared with the bovine gastrointestinal tract. Curli may therefore provide an important advantage during environmental persistence without being required for attachment to RSE cells.

The present observations are particularly consistent with our previous finding that curli can have a modulating rather than primary adhesive role in O157 adherence to bovine RSE cells (14). In that study, using curli variants and isogenic *csgA* mutants, we observed that removal of curli could actually increase adherence of certain O157 strains to RSE cells, in contrast to its more conventional role in attachment to human HEp-2 cells. This suggested that curli may physically temper or modify access of other bacterial surface structures to the bovine epithelial surface rather than serving as the dominant RSE adhesin (14). The current results extend that observation beyond a limited collection of O157 strains by showing that the same general lack of correspondence between curli phenotype and RSE adherence is evident in a larger collection containing O157, O26, and O111 STEC. In particular, the O26-6 isolate provides an important example of strong aggregative adherence despite limited environmental biofilm formation. Conversely, several curli-positive isolates did not display a correspondingly strong adherence phenotype. These observations make it unlikely that curli alone accounts for the strong RSE adherence phenotype. Instead, curli may contribute indirectly or conditionally allowing other adhesins or surface components to contribute to the RSE-binding phenotype.

The variation in Shiga toxin genotype and expression among the isolates provides another potential explanation for differences in adherence and warrants consideration. Shiga toxin has traditionally been regarded primarily as a cytotoxin responsible for host-cell injury, but evidence indicates that Stx2 can also influence bacterial colonization (18). Robinson et al. demonstrated that Stx2-producing O157:H7 exhibited enhanced adherence to HEp-2 cells and mouse intestinal colonization compared with an isogenic *stx_2_* mutant. Their experiments suggested that Stx2 enhanced adherence indirectly by increasing the availability of the eukaryotic receptor nucleolin, associated with intimin-mediated attachment rather than functioning as a conventional bacterial adhesin (18). At first glance, this mechanism could provide a possible explanation for the strong adherence observed among some Stx2-positive isolates in the present study. However, the current data do not support a major role for Stx2 in determining RSE adherence. O111 isolates lacking *stx_2_* displayed the same strong aggregative adherence phenotype as the O111 isolates carrying *stx_2_*. Furthermore, O157 and O111 isolates with different Shiga toxin profiles shared similar qualitative adherence patterns. Thus, while the published literature establishes that Stx2 can modulate epithelial adherence in particular O157 backgrounds, the present findings indicate that possession of *stx_2_* is neither necessary nor sufficient for bovine RSE cell adherence.

Our findings broadly address two stages of the STEC life cycle: environmental persistence and host colonization. Under environmental conditions, curli expression was associated with biofilm formation, particularly among O111 isolates. However, under host-associated conditions, biofilm production was greatly diminished. This suggests that the regulatory pathways controlling curli may favor persistence outside the host while other adhesins or surface structures become more important during host colonization. Previous studies have shown that relatively small changes in *csgD* regulation can produce substantial differences in curli expression and associated phenotypes in O157 (25). For instance, Uhlich et al. demonstrated that O157 curli variants with different *csgD* promoter states differed in their interaction with HEp-2 cells (25). These observations emphasize that curli regulation can affect bacterial-host interactions.

The observation in the present study that a few isolates retained detectable curli-expressing subpopulations despite an overall white phenotype is therefore potentially significant. Rather than viewing isolates simply as curli-positive or curli-negative, it may be more appropriate to consider curli as a regulated, heterogeneous phenotype that changes according to environmental conditions and/or bacterial physiological state. These findings have implications for understanding STEC persistence in cattle. The bovine recto-anal junction represents a major colonization site for O157 and other STEC, and the RSE assay provides a host-specific system for examining this interaction. The present results indicate that environmental persistence and host-cell attachment are not interchangeable phenotypes. The strong environmental biofilm phenotype of O111 may provide an advantage for persistence in cattle-associated environments, including surfaces and environmental niches outside the host. Once bacteria encounter the bovine epithelial environment, however, the regulatory conditions that promote curli-dependent biofilm formation may no longer function. Instead, adherence may depend on other surface structures that are expressed under host-associated conditions. The recent identification of multiple candidate accessory gene clusters associated with high RSE adherence (21, 26) further supports the concept that bovine epithelial attachment is likely to involve a complex repertoire of strain-specific factors.

Some limitations need to be factored in when interpreting results of the current study. First, the relationship between curli and RSE adherence was evaluated primarily through naturally occurring phenotypic variation rather than systematic genetic evaluation of the *csg* locus and its regulation. Consequently, a subtle contribution of curli cannot be fully excluded. Second, the biofilm assays measure bacterial accumulation on an abiotic surface, whereas the RSE assay measures interaction with a living bovine epithelial surface. Differences between the two phenotypes are therefore not completely unexpected. Future experiments involving targeted study of *csgA* or regulatory genes such as *csgD*, preferably in representative O157, O26, and O111 backgrounds, may provide better insights into a causal role of curli.

In conclusion, our results demonstrate that STEC adherence to bovine RSE cells is not simply a consequence of curli production or environmental biofilm-forming capacity. O157 and O111 isolates showed strong aggregative RSE adherence despite substantial variation in curli phenotype, whereas O26 isolates displayed greater heterogeneity in adherence. O111 isolates were particularly strong biofilm producers under environmental conditions, but this phenotype was substantially diminished under host-associated growth conditions. These findings support a model in which curli primarily contributes to environmental persistence and biofilm development, while RSE adherence is mediated predominantly by other, potentially strain- and serotype-specific surface factors. Curli may nevertheless modulate the bacterial surface and influence host interactions under particular conditions. The absence of a clear association between *stx_2_* and RSE adherence similarly suggests that Shiga toxin is unlikely to be the principal determinant of adherence in the STEC isolates tested, although a strain-specific or indirect role cannot be excluded. Taken together, the findings emphasize the importance of evaluating STEC adherence using host-relevant models and under physiologically relevant conditions rather than extrapolating directly from environmental biofilm phenotypes. The distinction between environmental persistence and host colonization may be particularly important for understanding how diverse STEC lineages persist in cattle and how they may be targeted for preharvest control.

## MATERIALS AND METHODS

### Bacterial strains and culture conditions

Thirty previously studied STEC isolates, ten each of O157:H7, O26:H11, and O111:H8 serotypes, were evaluated in this study (10). As listed in Table 3, the isolates originated from diverse sources (human, bovine, ground beef, and fly) and this diversity was represented among all serotypes studied. Bacterial freezer stocks were streaked individually on Luria-Bertani (LB) Lennox agar (Sigma-Aldrich, St. Louis, MO), incubated overnight at 37°C, and confirmed to be either O157 (*E. coli* O157 latex, Oxoid Diagnostic Reagents, Oxoid, Ltd., Hampshire, UK), O26, or O111 (*E. coli* non-O157 latex, Pro-Lab Diagnostics Inc., Round Rock, TX) *via* latex agglutination (27, 28). Latex-confirmed isolates were used in respective assays. Presence or absence of the virulence genes was verified in the DNA fingerprinting assay described below. Additional strains used as controls where needed included, (i) STEC O157 strain RM6067W, a non-curli producer (29), (ii) STEC O157 strain EDL933 (ATCC43895), a sequenced human clinical isolate (American Type Culture Collection, Manassas, Virginia; (30), and (iii) *E. coli* strain K-12, a well-characterized curli-producing bacterium (14).

### DNA fingerprint assay

The thirty isolates were typed using a technique called polymorphic amplified typing sequences (PATS) to determine their DNA fingerprint, as described in previous studies (31–34). Briefly, colony lysates were prepared for each isolate and specific primer pairs targeting 8 polymorphic *Xba*I- and 7 polymorphic *Avr*II-restriction enzyme sites as well as 4 virulence genes-*stx_1_* and *stx_2_* (Shiga toxins 1 and 2), *eae* (intimin-γ), and *hlyA* (hemolysin-A) were included (31, 32, 34–36). PCR amplicons of the *Avr*II-restriction enzyme site were purified with the QIAquick PCR purification kit (Qiagen, Valencia, CA) and digested with the *Avr*II restriction enzyme (New England Biolabs, Beverly, MA) to confirm the presence of the site. All reactions were gel electrophoresed on 3% GTG agarose gels stained with ethidium bromide. The presence or absence of *Xba*I and virulence genes amplicons was recorded as “1” or “0”, respectively. Additionally, the absence of an *Avr*II amplicon was recorded as “0”, presence of the restriction site with a single nucleotide polymorphism (SNP) as “1”, an intact restriction site as “2”, or “3” for a restriction site duplication (31, 32, 36).

### Immunoassay for Shiga toxin expression

An Stx-detecting enzyme-linked immunosorbent assay (ELISA) was used to evaluate the presence or absence of Shiga toxins in the thirty isolates and STEC O157 strain RM6067W, with *E. coli* strains K-12 and STEC O157 strain EDL933 (ATCC 43895) serving as bacterial negative and positive controls, respectively. The Premier® EHEC kit (Meridian Bioscience, Cincinnati, OH; https://www.meridianbioscience.com/diagnostics/disease-areas/gastrointestinal/e-coli/premier-ehec/), which detects both Shiga toxins I and II, was utilized per the manufacturer’s instructions. Briefly, the STEC strains were streaked on fresh LB agar and incubated overnight at 37°C. Individual colonies were inoculated in 5 mL Difco GN broth, Hanja (BD Biosciences, Franklin Lakes, NJ) and incubated static overnight at 37°C. Overnight cultures (50μL) were diluted in sample diluent (Premier® EHEC kit) and 100 µL was used to set up the ELISA, in duplicate. Plates were subsequently read at 450 nm on a Synergy HT microplate reader (BioTek Instruments, Winooski, VT) with medium shaking for 5 sec. Samples were considered positive for the presence of Shiga toxins when the OD_450nm_ was ≥0.180 (Premier® EHEC kit). The average post-EHEC-ELISA absorbances were further used to derive a toxin gene / expression score as follows: isolates with the presence of Stx1-only were given a “1”, isolates with Stx2-only a “2”, and isolates with both toxins a “3”; for the expression score-isolates with post-assay absorbances between 0.19-1.5 were designated as “+”, isolates between 1.6-2.5 a “++”, and isolates between 2.6-3.5 as “+++”.

### Recto-anal junction squamous epithelial (RSE) cell adherence

Adherence patterns of the thirty STEC isolates to RSE cells were evaluated in an *in vitro* assay, conducted as previously reported (14, 17, 19). Briefly, RSE cells were diluted to a final concentration of 10^5^ cells/mL in DMEM-NG and mixed with bacterial isolates at a bacteria:cell ratio of 10:1. The mixture was incubated for 4 h at 37°C with shaking at 110 rpm, pelleted, and washed in double-distilled water (dH_2_O). The final pellet was resuspended in 100 µL of dH_2_O and 2 µL drops were placed on Polysine slides (Thermo Scientific), dried, and fixed. Slides were IF stained with antibodies specific to RSE cell cytokeratins and to the O157 antigen or *E. coli* (14, 16, 17). Bacterial adherence patterns on the RSE cells were qualitatively recorded as diffuse, aggregative, or non-adherent and quantitatively recorded as RSE cell percentages with or without adherent bacteria (20). Adherence was further categorized as: hyper-adherent when more than 50% of RSE cells had 10 adherent bacteria; moderate when ≤50% had 5-10 adherent bacteria; and non-adherent when ˂50% of RSE cells had 1-5 adherent bacteria. RSE cells without bacteria were included as a negative control and the assay was conducted in two biological replicates with eight technical replicates each.

### Biofilm and curli formation assays

STEC strains were grown in Luria-Bertani broth with no salt (LB-NS) overnight at 26°C. Each overnight was sub-cultured onto fresh Congo Red Indicator (CRI) agar (LB-NS agar, 20 mg/mL Congo red dye, 5 mg/mL Coomassie brilliant blue) and diluted in fresh LB-NS broth to a concentration of 10^5^ CFU/mL, of which 125 μL was aliquoted into 3 wells on Corning® round-bottom 96-well polystyrene plates (Corning, Inc., Corning, NY), in triplicate. Plates were incubated static at 26°C for 2 days (48 h) in a plastic bag with a moist tissue for humidity. The planktonic cells were removed post-incubation, and the wells were rinsed twice with 150 μL sterile distilled water, heat fixed at 80°C for 30 min, and stained with 150 μL 0.1% crystal violet at room temperature for 30 min. After staining the biofilm-associated bacterial cells, the dye was removed, and the wells were washed four times with sterile distilled water and the bound crystal violet was solubilized in 150 μL 33% acetic acid for 10 min. The absorbance was read at 570 nm with a microplate reader (SpectraMax 190; Molecular Devices, Sunnyvale, CA or Synergy HT; BioTek Instruments).

Mixed CRI variants were isolated *via* sub-culture, stocked at -80°C, and evaluated separately using the same methodology as described above. For strains or mixed CRI variants that were Congo Red-positive (CR+) but did not yield a biofilm in the 48 h plate assay, a 5 d (120 h) LB-NS tube biofilm formation assay was conducted. Briefly, 5ml diluted overnight cultures (10^5^ CFU/mL) were dispensed into a 16×150 mm borosilicate glass tube, in triplicate, and incubated static at 26°C for 5 days. The planktonic cells were then removed, and the tubes were rinsed with sterile distilled water. The bound dye was solubilized in 6 mL 33% acetic acid for 10 min and the absorbance was measured as described above.

Furthermore, eight of the thirty isolates (O157-1W, O157-9P, O26-6P, O26-9W, O26-10R, O111-2R, O111-3W, and O111-8R) were selected for additional analysis. Strains were evaluated in 2 d plate biofilm assays in Dulbecco’s modified Eagle’s medium (DMEM) with low glucose (DMEM-LG; Invitrogen, Carlsbad, CA) at both 26°C, 37°C and 39°C in a similar fashion as described above. The isolates were also subcultured on CRI plates under the same temperature conditions.

Each biofilm assay included controls of media-only and *E. coli* strain K-12, STEC O157 strain EDL933 (ATCC 43895), and STEC O157 strain RM6067W. The average adjusted biofilm OD_570nm_ was calculated by subtracting from that of the media-only and the resulting value was used for scoring. Average adjusted OD_570nm_ less than 0.5 were considered negative for biofilm formation, “+” for 0.6-1.0, “++” for 1.1-2.0, “+++” for 2.1-3.0, >3.1 as “++++”, and “T” if a biofilm formed during a tube assay. In addition, CRI phenotypes were recorded based on colony pigmentation following 48 h incubation at 26°C as follows: i) curli-deficient white colonies (C-), ii) curli-producing smooth, pink colonies (C+), iii) curli-producing red colonies (C++), and iv) curli-producing super-red colonies (C+++).

### Immunofluorescent (IF) microscopy

Seventeen CRI curli-negative isolates were evaluated for curli production *via* IF staining (O157 isolates 1W, 2W, 3W, 4W, 5W, 6W, 7W, 8W, 10W; O26 isolates 2W, 5W, 9W, 10W; and O111 isolates 3W, 5W, 6W, 9W) along with the aforementioned control strains. After a 5 d tube incubation in LB-NS at 26°C for biofilm formation (as described above), planktonic cells were removed, and biofilms, if any, were gently scraped off the sides of the tubes with 1 mL sterile phosphate-buffered saline (PBS). Biofilm cell suspensions and 1 mL of corresponding planktonic cells were centrifuged, and the pellets were washed in an additional 1 mL sterile PBS. Pellets were resuspended in 100 µL sterile distilled water and smears were prepared on Polysine slides (Thermo Scientific, Rockford, IL) for IF-curli staining as previously described (14). Slides were fixed with ethanol and blocked with 5% normal goat serum (NGS; Vector Laboratories, Burlingame, CA)-PBS. Slides were then incubated with absorbed rabbit anti-curlin diluted 1:20 in PBS for 1 h at room temperature, washed in PBS, and incubated with secondary antibodies diluted in PBS for 1 h at room temperature. The secondary antibodies consisted of AlexaFluor 594-labeled goat anti-rabbit IgG [H+L; F(ab’)_2_ fragment] (Invitrogen) targeting anti-curlin primary antibody and the fluorescein isothiocyanate (FITC)-labeled goat anti-O157 or anti-*E. coli* (KPL, Gaithersburg, MD) antibodies labelling O157 or *E. coli*, respectively. Slides were washed with PBS after secondary antibody incubation with a final wash in distilled water and air-dried. Coverslips were added with Prolong Glass antifade reagent (Invitrogen) and analyzed on a Nikon Eclipse E800 fluorescence microscope (Nikon Instruments, Inc., Elgin, IL) with appropriate filters and digital imaging capabilities.

### Statistics

All analyses for the EHEC-ELISA, biofilm, and RSE adherence assay data were conducted in GraphPad Prism (version 8.4.3, GraphPad Software, San Diego, CA). Descriptive statistics and standard deviation and/or standard error of the mean were used for the absorbance data (EHEC-ELISA OD_450nm_ data and biofilm OD_570nm_), and for RSE bacterial adherence counts, respectively. The adherence assay data and the biofilm post-assay average adjusted OD_570nm_ values were evaluated with unpaired *t*-tests and one-way ANOVAs. A significance level of *p* <0.05 was considered statistically significant.

## ACKNOWLEDGEMENT

Excellent technical support provided by *Late* Bryan K. Wheeler, Biological Scientific Technician, is gratefully acknowledged. We thank Drs. Terrance Arthur, Rong Wang, and James Bono at the Roman L. Hruska U.S. Meat Animal Research Center/ARS/USDA for the 30 STEC strains, and Dr. Michelle Q. Carter at the Western Regional Research Center/ARS/USDA for the O157 strain RM6067W used in this study. The support provided by the NADC Animal Resource Unit with the bovine RAJ tissues collection at necropsies is much appreciated. This work was supported by USDA-ARS CRIS projects 5030-32000-225-00D (ITK). HM was supported by the USDA/1890 Scholarship program. ENB, KD were supported by an appointment to the ARS Research Participation Program administered by ORISE through an interagency agreement between the U.S. DOE and the USDA. ORISE is managed by ORAU under DOE contract number DE-SC0014664.

## Disclaimer

Mention of trade names or commercial products in this article is solely for the purpose of providing specific information and does not imply recommendation or endorsement by the U.S. Department of Agriculture. USDA is an equal opportunity provider and employer.

## SUPPLEMENTAL MATERIALS

**Table S1.** PATS profiles of the thirty STEC isolates.

**Table S2.** Adherence characteristics of STEC isolates on RSE cells.

**Table S3.** Summary of 26°C LB-NS biofilm 2 d and 5 d assay data for all original thirty STEC isolates and controls.

**Table S4.** Summary of biofilm data for select STEC isolates and controls. 26°C LB-NS biofilm 2 d and 5 d data for all original thirty STEC isolates

**Figure S1.** Representative adherence patterns of Control strains (Panel A), STEC O157 (Panel B), O26 (Panel C) and O111 (Panel D) isolates on RSE cells. The immunofluorescence stained slides are shown at 40x magnification. Bacteria have green fluorescence, RSE cells’ cytokeratins have orange-red fluorescence, and the nuclei have blue fluorescence. Numbers represent each different isolate per serotype.

**Figure S2.** Representative images of immunofluorescence-labeled bacteria and curli are shown along with a descriptive table. Images are shown at 100x objective magnification. Bacterial cells have green fluorescence and the enveloping curli have orange-red fluorescence.

**Figure S3.** Summary of curli phenotype observed on CRI plates for the control strains and select STEC isolates. Images for variant phenotypes at different incubation temperatures are shown separately. Putative curli-producers made red or smooth, pink colonies while curli non-producers were colorless/white as indicated in the inserted table.

